# The HCoV-HKU1 N-terminal domain binds a wide range of 9-*O*-acetylated sialic acids presented on different glycan cores

**DOI:** 10.1101/2024.05.24.595699

**Authors:** Ilhan Tomris, Anne Kimpel, Ruonan Liang, Roosmarijn van der Woude, Geert-Jan Boons, Zeshi Li, Robert P. de Vries

## Abstract

Coronaviruses recognize a wide array of protein and glycan receptors using the S1 subunit of the spike (S) glycoprotein. The S1 subunit contains two functional domains: the N-terminal (S1-NTD) and C-terminal (S1-CTD). The S1-NTD of SARS-CoV-2, MERS-CoV, and HCoV-HKU1 possess an evolutionarily conserved glycan binding cleft that facilitates weak interactions with sialic acids on cell surfaces. HCoV-HKU1 employs 9-*O*-acetylated α2-8-linked disialylated structures for initial binding, followed by TMPRSS2 receptor binding and virus-cell fusion. Here, we demonstrate that HCoV-HKU1 NTD has a broader receptor binding repertoire than previously recognized. We presented HCoV-HKU1 NTD Fc chimeras on a nanoparticle system to mimic the densely decorated surface of HCoV-HKU1. These proteins were expressed by HEK293S GNTI^-^ cells, generating species carrying Man-5 structures, often observed near the receptor binding site of CoVs. This multivalent presentation of high-mannose-containing NTD proteins revealed a much broader receptor binding profile compared to its fully glycosylated counterpart. Using glycan microarrays, we observed that 9-*O*-acetylated α2-3 linked sialylated LacNAc structures are also bound, comparable to OC43 NTD, suggesting an evolutionarily conserved glycan-binding modality. Further characterization of receptor specificity indicated promiscuous binding towards 9-*O*-acetylated sialoglycans, independent of the glycan core (glycolipids, *N-* or *O*-glycans). We demonstrate that HCoV-HKU1 may employ additional sialoglycan receptors to trigger conformational changes in the spike glycoprotein to expose the S1-CTD for proteinaceous receptor binding. (218)

## Introduction

Coronaviruses (CoVs) cause respiratory and digestive disorders, manifesting in various species, including dogs, cats, cattle, and other animals [1]. These CoVs are separated into four genera based on their genomic structure and phylogenetic branching: Alpha (α), Beta (β), Gamma (γ), and Delta (δ), with many unidentified CoVs circulating in the natural reservoir (bats and rodents) [2]. Several α- and β-genus CoVs (Table S1) have crossed the human barrier and may cause mild to severe respiratory tract infections. These human coronaviruses (HCoVs) differ in receptor recognition, virus internalization pathways, and accessory genes. Receptor binding is driven by the spike (S) glycoprotein, which is displayed on the viral membrane and consists of two functional subunits: S1 and S2. The S1 facilitates cellular receptor interaction and is divided into the N-terminal domain (S1-NTD) and the C-terminal domain (S1-CTD). The binding of CoVs to specific receptors defines tropism, and both the S1-NTD and S1-CTD may function as the receptor binding domain (RBD) [1, 3]. The S1-NTDs of CoVs contain an evolutionarily conserved galectin fold that functions as a (sialo)glycan-binding viral lectin [4], while the S1-CTD interacts with proteinaceous receptors. Following conformational rearrangement of S2, initiated by S1-receptor binding, membrane fusion is triggered, and viral entry occurs through the plasma membrane or endosomal pathway [5].

The sugar layer covering eukaryotic cells, the glycocalyx, serves as a barrier against pathogens. Viral attachment is a crucial and dynamic process in host-cell infections, and several pathogens use sialoglycans as an initial attachment point [6], an important determinant for tropism and defining the host range [7, 8]. Sialylated glycans are employed by several (human) CoVs, albeit further modifications of the sialic acid (SIA) influence receptor attachment [9, 10]. Glycan structures with terminal SIA (α2–3, α2–6, and α2–8-linked) exist in several modified forms [11]. These commonly occur on carbon 4, 5, 7, 8, and 9 positions, with regulation being dependent on physiological conditions and animal species [7, 9]. The *O*-acetyl modification of SIA is introduced in the Golgi apparatus by SIA-specific *O*-acetyltransferases (SOATs), with CAS1 domain containing 1 (CASD1) being the only SOAT identified so far [12]. The exact role of *O*-acetylation is not well-defined, and the abundance of this modification appears to differ per tissue [7, 8]. SARS-CoV-2, Bovine CoV (BCoV), HCoV-OC43, and HCoV-HKU1 engage 9-*O*-acetylated SIAs using their S1-NTD [13-16]. Interestingly, although these CoVs belong to the β-CoV genus, these viruses do not cluster in the corresponding subgenera. SARS-CoV-2 belongs to the Sarbecovirus subgenus, while BCoV, HCoV-OC43, and HCoV-HKU1 are Embecoviruses [17], suggesting that 9-*O*-acetylated SIA recognition is an evolutionarily conserved property between CoVs [13, 16, 18]. HCoV-OC43 and HCoV-HKU1 show high sequence similarity and recognize the same receptor; however, these viruses were introduced and adapted to the human population independently [19, 20]. HCoV-OC43 and HCoV-HKU1 S1-NTD both recognize 9-*O*-acetylated α2-8-linked disialic acid, whilst HCoV-OC43 displays promiscuity towards 9-*O*-acetylated α2-3- and α2-6-linked SIA [10, 16].

Virus-glycan interactions are inherently of low binding affinity, requiring multivalent display for receptor characterization. Multivalent effects can be attained with precomplexation using antibodies [21, 22], virus-like particles (VLPs) and nanoparticles (NPs) highly resemble pathogens by presenting antigenic epitopes in a highly dense and ordered array, enabling multiple binding events [23, 24]. In immunization and vaccine development, these multivalent molecules are favored compared to monovalent equivalents [25]. Similarly, to increase the sensitivity through an increase of avidity, HCoV-HKU1 S1-NTD was presented on a self-assembling 60-mer to probe for sialoglycan binding and receptor specificity. Here, we elaborate that multivalent presentation enabled HCoV-HKU1 S1-NTD to bind to the more abundant 9-*O*-acetylated α2-3 linked SIA containing epitopes. Although HCoV-HKU1 and HCoV-OC43 are separated by considerable phylogenetic distance, glycan binding of HCoV-HKU1 is thus nearly identical to that of HCoV-OC43 [10, 15], indicating that the mode of receptor-binding is conserved between Embecoviruses [15].

## Results

### Glycosylation and multivalency in virus-receptor interaction

Several naturally occurring nanoparticle platforms enable mimicry of pathogens, by displaying multiple viral envelope proteins in a spherical form [26-28]. Here, we employ a self-assembling NP scaffold based on Lumazine Synthase (LS) from *A. aeolicus* fused with domain B of protein A (pA) from *S. aureus* [28, 29]. The pA part from the pA-LS scaffold (60-mer) enables high-affinity interaction with antibody Fc regions; thus, to present HCoV-HKU1 NTD on the NP scaffold, HCoV-HKU1 NTD Fc chimeras were expressed and combined with pA-LS (Fig. 1A). HCoV-HKU1 NTD, expressed in 293T and used as a lectin only, recognizes the 9-*O*-acetylated α2-8-linked disialic acid (#**12**) as previously described (Fig. 1B and Table S2) [10, 15]. The same binding pattern was observed when this protein was precomplexed using antibodies or as a nanoparticle. Expressing sialic acid-binding proteins with complete glycosylation may result in aggregation and dim binding signals, as previously observed for the hemagglutinin protein of the influenza A virus [30, 31]. To overcome the negative effect of complex glycosylation in receptor binding, we employed *N*-acetylglucosaminyltransferase I (GnTI) deficient cells [32] to assess the possible broader receptor specificities of HCoV-HKU1 NTD. Non- and pre-complexed NTD Fc chimeras did not recognize additional structures; however, with a multivalent presentation using pA-LS NP, binding to a 9-*O*-acetylated α2-3 Sia LacNAc structure was observed (Fig. 1B and Table S1). The expression of HCoV-HKU1 NTD in GnTI deficient cells and multivalent presentation on pA-LS NP enabled characterization of previously undetected receptor-binding specificity to an abundant terminal epitope.

**Fig. 1.**
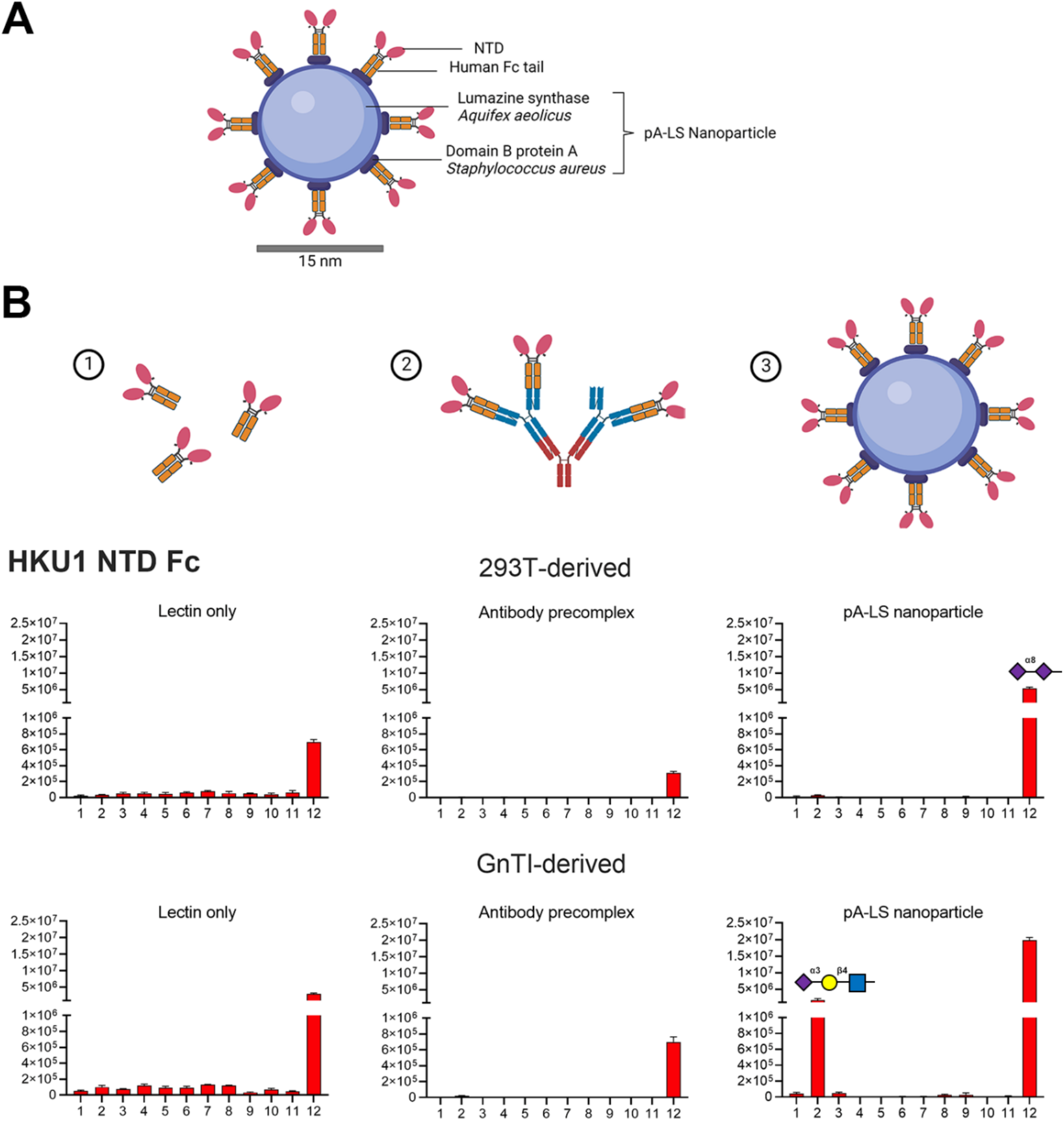
Multimerization-dependent receptor probing. **(A)** (1) HKU1 NTD Fc, (2) precomplexation using antibodies, and (3) HKU1 NTD Fc chimera presented on a 60-mer pA-LS nanoparticle. **(B)** HKU1 NTD Fc derived from HEK293T cells displayed receptor specificity towards 9-*O*-acetylated α2-8 linked disialylated structures as lectin only, precomplexed with antibodies or presented on a NP. HKU1 NTD Fc derived from HEK293S GnTI-deficient cells recognize 9-*O*-acetylated α2-8-linked disialylated structures when used as lectin only or antibody precomplexed, an additional α2-3-linked SIA LacNAc structure is bound when HKU1 NTD is presented on a NP.

### HKU1 recognizes O-acetylated SIA on different glycoconjugates

Previously, disialylated gangliosides were indicated as a cellular receptor for HCoV-HKU1; however, ST8Sia1 was required to facilitate binding towards non-susceptible HEK293T cells [10]. The ST8Sia1 enzyme transfers SIA to an α2-3-linked SIA in GM3, thus generating α2-8-linked disialylated GD3. SIAs with an α2-8-linkage are formed through eight different α2-8-sialyltransferases (ST8Sia1-8) [33] and the acceptor substrate (i.e., glycolipid, *N*- or *O*-glycan) varies per ST8Sia. Thus, the contribution of *N*- and *O*-glycans in receptor binding was not assessed due to the exclusion of ST8Sias that transfer SIA to these glycan cores.

To assess the contribution of *N*- and *O*-glycans in addition to glycolipids, the rhabdomyosarcoma (RD) cell line was employed as this cell line is susceptible for live HCoV-HKU1 virus [34]. HCoV-HKU1 NTD was expressed as a trimer using a GCN4 trimerization domain (HCoV-HKU1 NTD tri) [21], or as an Fc chimera (HCoV-HKU1 NTD Fc). For both trimeric and Fc HCoV-HKU1 NTD variants, binding was observed on RD cells (Fig. 2). Fluorescence intensity of trimeric NTD in comparison to NTD Fc appeared to be higher, related to additional receptor binding sites in a trimer versus a dimer.

**Fig. 2.**
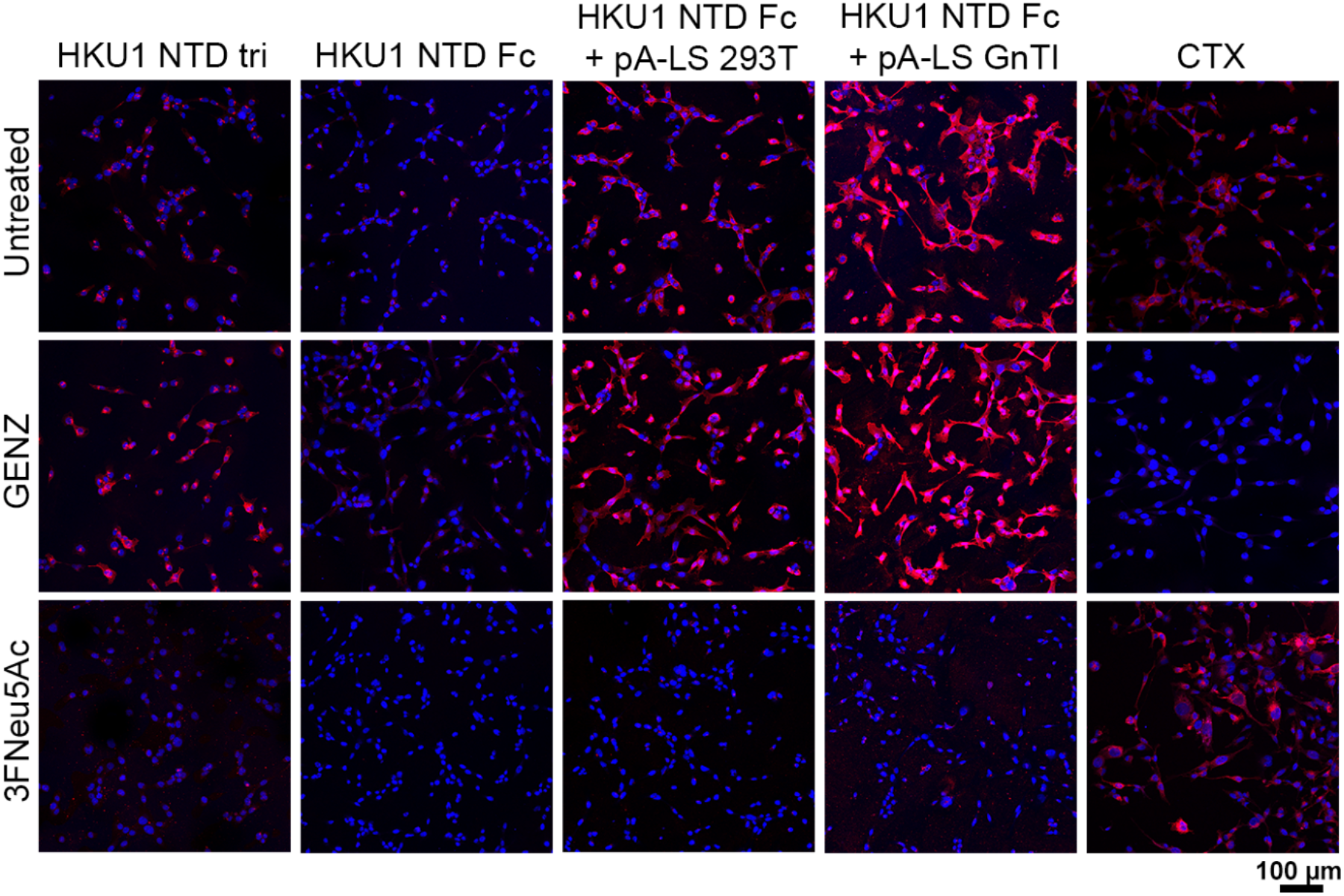
Glycolipid-independent receptor specificity. Trimeric HKU1 NTD, HKU1 NTD Fc, and HKU1 NTD Fc + pA-LS (GnTI-derived and 293T-derived) binding to RD cells. Utilization of a glycolipid inhibitor (GENZ-123346) did not result in a reduction of fluorescence intensity for HKU1 NTD. Binding of HKU1 NTD was abrogated when cells were treated with 3FNeu5Ac. CTX binding was completely abrogated after the addition GENZ, whilst 3FNeu5Ac treatment did not influence binding.

Multivalent presentation of HCoV-HKU1 NTD Fc on pA-LS NP increased the fluorescence intensity. When comparing HEK293T- and HEK293S GnTI^-^-derived proteins, a notable increase in signal was observed for the GnTI^-^-derived HCoV-HKU1 NTD. This boost in binding is probably related to the observation that an additional 9-*O*-acetylated α2-3-linked Sia LacNAc structure is bound. Treatment of RD cells with a glycolipid inhibitor (GENZ-123346) was performed to characterize the contribution of sialoglycolipids in receptor binding (Fig. 2). Inhibition of glycolipid synthesis was verified using Cholera toxin (CTX), but no decrease of binding for HCoV-HKU1 NTD was observed [10, 16]; clearly, *N*- and *O*-sialoglycans can compensate for glycolipid inhibition. To verify that SIAs are indeed the determinant for receptor interaction, a sialyltransferase inhibitor (3FNeu5Ac) was employed, resulting in complete abrogation of HCoV-HKU1 NTD binding. In conclusion, we demonstrate glycolipid-independent receptor binding of HCoV-HKU1 NTD. The contribution of different glycoconjugates presented on cell surfaces needs to be taken into consideration to define receptor binding specificity profiles of viruses.

### Redundancy in receptor recognition for HKU1 coronavirus

The sialoglycome on cell surfaces consists of glycolipids, *N*- and *O*-glycans. Glycolipid-independent binding of HCoV-HKU1 NTD warranted further characterization of *N*- and *O*-glycan contribution in receptor recognition. The synthesis of complex *N*-glycans was inhibited with swainsonine; this compound blocks further processing of high-mannose *N*-glycans by Golgi α-mannosidase II. Inhibition of *N*-glycans elaborated that binding of HCoV-HKU1 NTD, as trimer and (multivalent) Fc chimera, was not dependent on *N-*glycosylation (Fig. 3A). Phytohemagglutinin-L (PHA-L) lectin was utilized to assess *N*-glycan inhibition; swainsonine treatment indeed abrogated binding, while this was not the case with Ac5GalNTGc inhibition (Fig. 3B). This Ac5GalNTGc inhibitor was employed to study the contribution of *O-*glycans as it blocks the transfer of galactose by Core 1 β1,3-galactosyltransferase (C1GalT1) to GalNAcα1-Ser/Thr (Tn-antigen), resulting in trimmed *O*-glycans [35]. Although overall fluorescence intensity was diminished, the trimeric NTD, precomplexed Fc chimera, and Fc + pA-LS still bound, indicating binding to non-*O*-glycosylated structures (Fig. 3A). Verification of *O*-glycan trimming using Ac5GalNTGc was performed using peanut agglutinin (PNA) and Vicia villosa lectin (VVL): these lectins recognize the Tn-antigen and an increase in fluorescence intensity indicates *O*-glycan inhibition/trimming. The addition of the inhibitor did not improve the binding of PNA and VVL to RD cells (Fig. 3C), probably due to aberrant sialylation as is common in cancerous cells, like the RD cells employed here, which blocks the binding of these lectins [36]. The influence of SIA on binding was assessed by utilizing 3FNeu5Ac; the fluorescence signal increased for both PNA and VVL, albeit with a more significantly for PNA (Fig. 3C). To confirm *O*-glycan trimming, cells were treated with 3FNeu5Ac + Ac5GalNTGc. An increase of intensity concerning 3FNeu5Ac treated cells was observed for PNA and VVL; however, this was most pronounced for VVL.

**Fig. 3.**
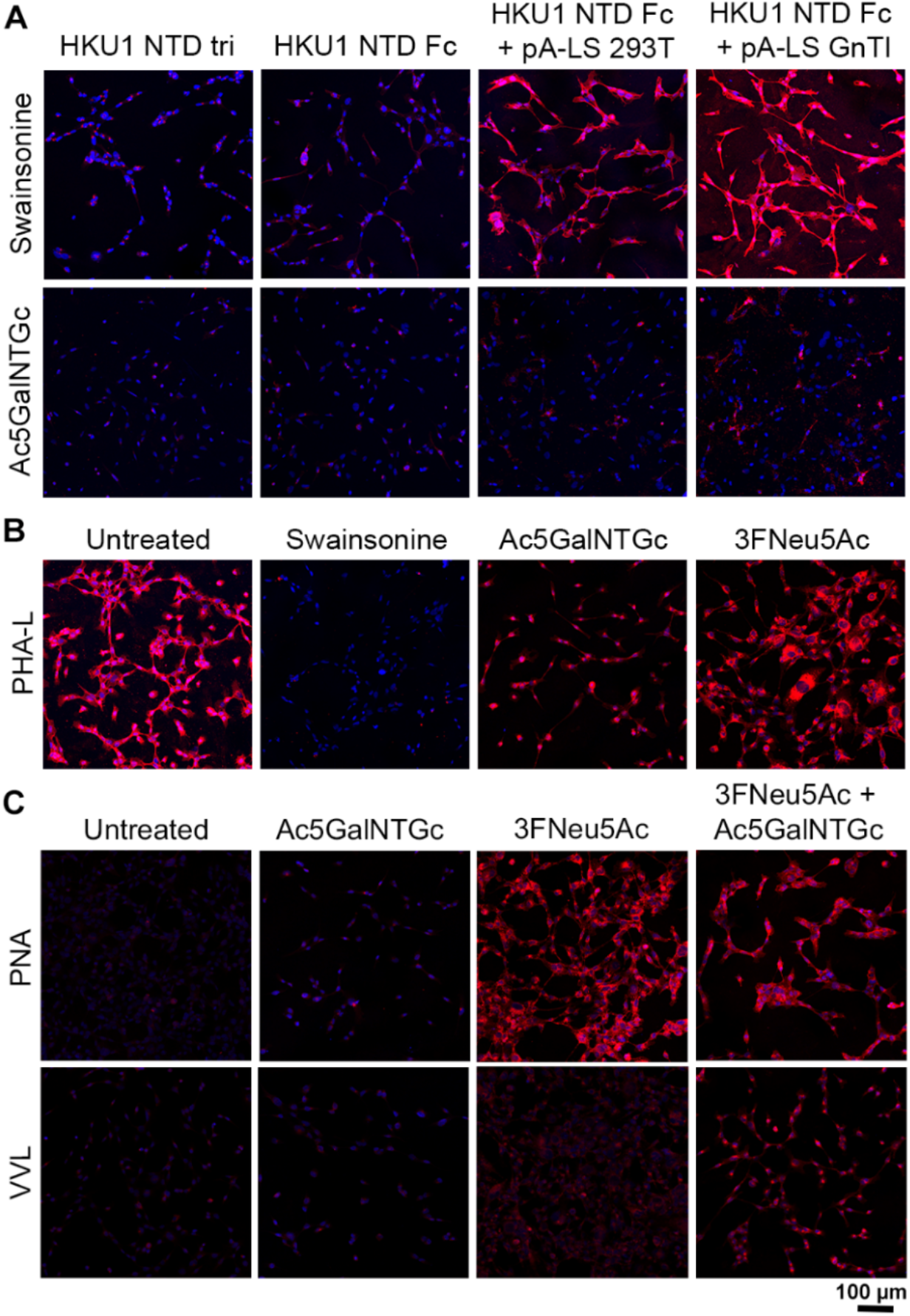
Receptor recognition is independent of the glycan core. **(A)** N-glycan-independent receptor binding for trimeric HKU1 NTD, HKU1 NTD Fc and HKU1 NTD Fc + pA-LS (GnTI-derived and 293-derived), receptor binding was not influenced by the addition of a complex *N*-glycan inhibitor (swainsonine). Fluorescent signal was significantly reduced after the use of an *O*-glycan inhibitor (Ac5GalNTGc), albeit spots with high fluorescence intensity were still observed. **(B)** Complex N-glycan synthesis inhibition was verified with PHA-L lectin, a reduction of fluorescence intensity was visualized with the use of swainsonine, whilst Ac5GalNTGc and 3Fneu5Ac did not influence PHA-L binding. **(C)** PNA and VVL lectins used as controls for *O*-glycan truncation after Ac5GalNTGc treatment.

We further characterized core glycan independent binding using flow cytometry. We first verified the inhibition of glycolipids, *N*- and *O*-glycans using CTX, PHA-L, PNA, and VVL lectins (Fig. 4A). Interestingly, Ac5GalNTGc treatment significantly increased (p < 0.05) CTX binding to RD cells, suggesting that glycolipid synthesis was upregulated to compensate for *O*-glycan inhibition. Similar to confocal imaging data, trimeric HCoV-HKU1 NTD and NTD Fc + pA-LS binding was abolished by 3FNeu5Ac treatment (Fig. 4B). Additionally, treatment with GENZ, swainsonine, and Ac5GalNTGc did not display any dependence on the inhibited glycan core. Further characterization with a combination of inhibitors, such as swainsonine + GENZ, swainsonine + Ac5GalNTGc, and Ac5GalNTGc + GENZ, to specifically isolate the individual glycan cores was performed. Despite inhibiting a significant portion of the sialoglycome, HCoV-HKU1 NTD binding was maintained and is thus fully redundant. This suggests that 9-*O*-acetylated structures are presented on various core glycans and that promiscuous binding employed by HCoV-HKU1 could improve its infectivity.

**Fig. 4.**
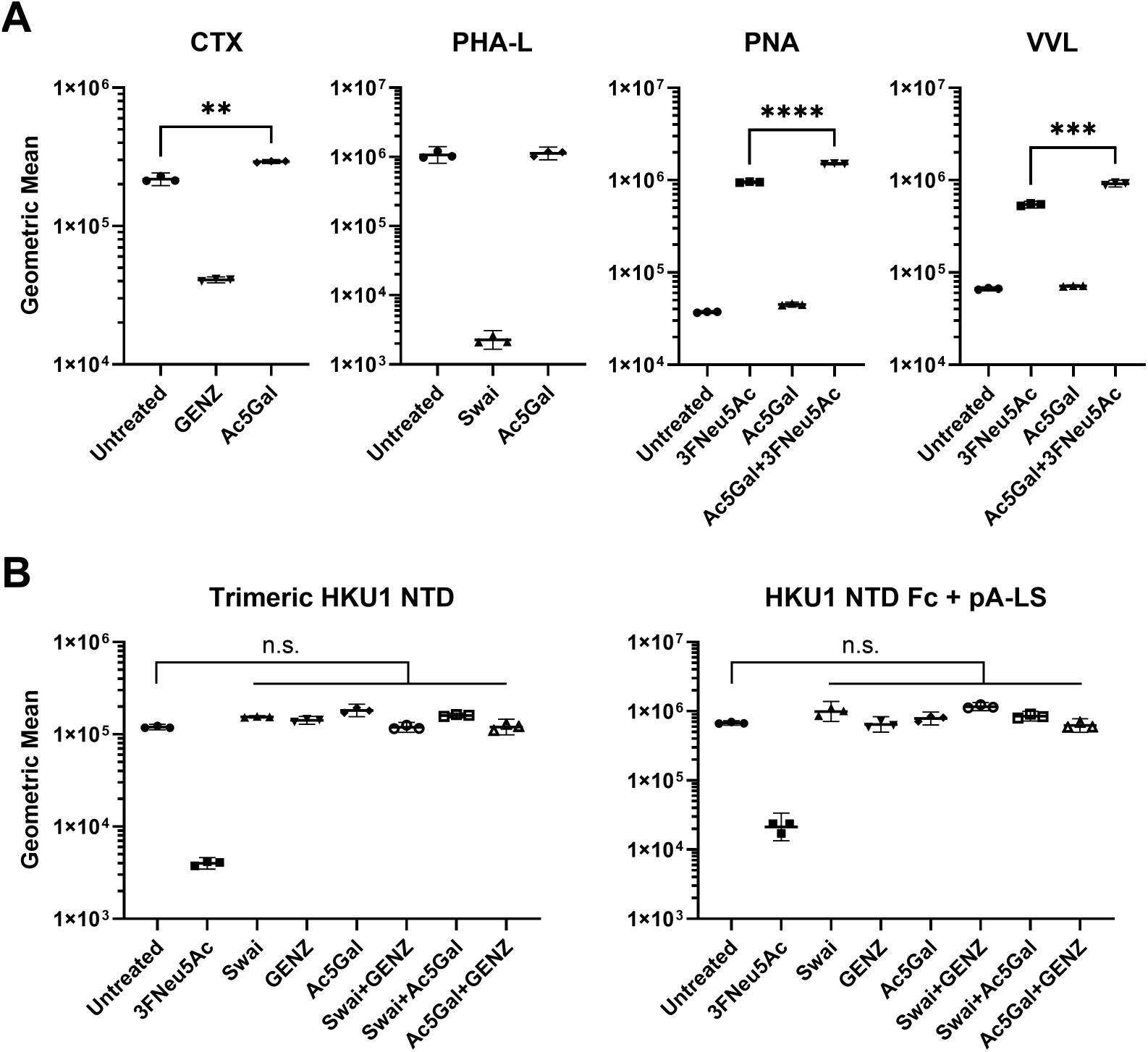
Redundant glycan receptor recognition of HKU1 NTD. Inhibition of glycolipid, *N*- and *O*-glycan truncation on RD cells verified using flow cytometry. **(A)** CTX and PHA-L fluorescence signal decreased after inhibitor treatment. Desialylation with 3FNeu5Ac increased PNA and VVL binding, hereafter combination of 3FNeu5Ac with Ac5GalNTGc enabled observation and confirmation of *O*-glycan truncation. **(B)** Receptor recognition of trimeric HKU1 NTD and HKU1 NTD Fc presented on pA-LS NP was only influenced by desialylation. ** = p < 0.05, **** = p < 0.0001, n.s. = not significant. Swai, swainsonine; Ac5Gal, Ac5GalNTGc.

### Mucins as a potential first point of contact for HCoV-HKU1

Mucus forms a protective layer of the airway epithelium and serves as the first line of defense [37, 38]. The major macromolecular components of mucus are heavily *O*-glycosylated and *O*-acetylated glycoproteins [8, 37, 39, 40], known as mucins. HCoV-HKU1 NTD engages 9-*O*-acetylated SIAs, independent from the glycan core. Since mucins play a protective role in the airways against pathogens, we aimed to study the interaction with bovine submaxillary mucin (BSM) as it is heavily modified with *O*-acetylated SIA. Fetuin was used as a control; this 48 kDa molecule does not contain *O-*acetylated structures but is heavily sialylated. Trimeric HCoV-HKU1 NTD and Fc chimeras derived from HEK293T or HEK293S GnTI-deficient cells were analyzed for binding to both substrates. HCoV-HKU1 NTD Fc precomplexed with antibodies or presented on pA-LS NP displayed similar binding properties to BSM (Fig. 5A), albeit stronger for the NP (Table S3). In contrast, no binding was observed for fetuin, as expected as this substrate does not display 9-*O*-acetylated structures (Fig. S1). The trimeric NTD failed to bind both BSM and fetuin (Fig. 5B and Fig. S1). The difference in binding based on the glycosylation state was pronounced, indicating the necessity of utilizing proteins derived from HEK293S GnTI-deficient cells rather than HEK293T-derived proteins with complex glycosylation. The presence of α2-3- and α2-6-SIA was confirmed with the use of MAL-II (α2-3 SIA, BSM) and SNA lectins (α2-6 SIA, BSM) (Fig. 5B and Fig. S1). Thus, BSM contains α2-3- and α2-6-linked sialylation and *O*-acetylation whilst lacking α2-8 disialylated SIAs. Importantly, BSM does not contain disialylated structures [41] Thus the presence of 9-*O*-acetylation is the driving factor for glycan binding, independent of the glycan core.

**Fig. 5.**
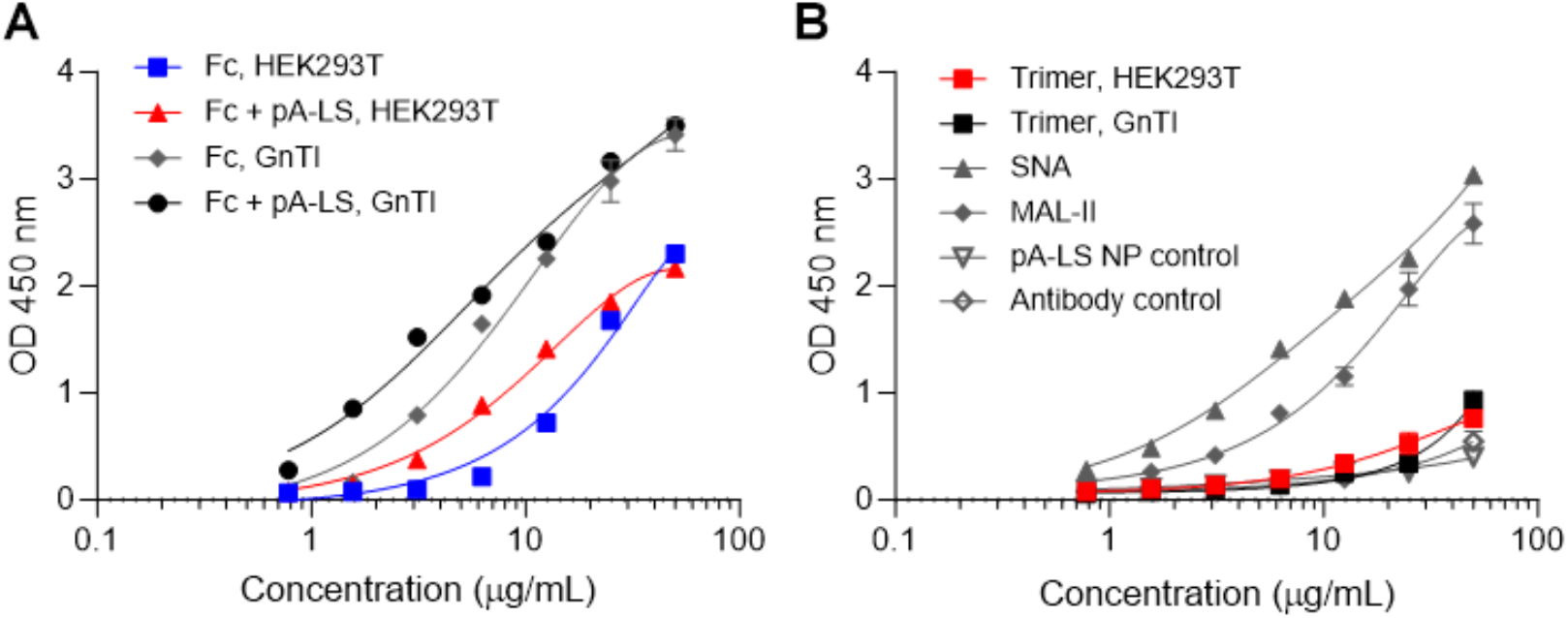
Virolectin binding is α2-8-linked-independent. **(A)** HEK293S GnTI-derived HKU1 NTD Fc and HKU1 NTD Fc + pA-LS displayed increased binding to BSM in comparison to their 293T-derived equivalents. **(B)** Presence of α2-3-SIA on BSM was confirmed by utilizing MAL-II, while SNA binding confirmed the presence of α2-6-SIA. Trimeric HKU1 NTD (GnTI- and 293-derived), pA-LS NP, and antibody control did not bind BSM.

## Discussion

Here we demonstrate that multivalency and the glycosylation state influence glycan binding properties of the HCoV-HKU1 NTD protein; enabling the observation of binding 9-*O*-acetylated α2-3 linked Sia LacNAc binding. Furthermore, additional biochemical experiments were performed to assess the contribution of glycolipids in NTD-glycan binding, as α2-8 linked SIA is often a terminal epitope of such glycans. Inhibition of glycolipid synthesis did not influence the receptor-binding properties of HCoV-HKU1 NTD, indicating the importance of *N-* and *O-*glycans. Similarly, the inhibition of *N-* or *O-*glycans did not elucidate preferential binding to specific glycan cores (glycolipid, *N-* or *O-*glycan). The utilization of Ac5GalNTGc as an *O-*glycan inhibitor reduced the overall fluorescence intensity using confocal microscopy, however, intense fluorescence was still present in certain locations on the cell, resulting in robust signal using flow cytometry. We conclude that HCoV-HKU1 NTD binding to 9-*O-*acetylated SIAs is independent of the glycan core and recognition of non-disialylated structures, warranting further investigation. We then employed BSM as a ligand, which is a heavily *O-*glycosylated and *O-*acetylated glycoprotein that does not contain α2-8-linked disialylated structures [8, 37, 39-41]. HCoV-HKU1 NTD appeared to interact efficiently with the ligands displayed on BSM, containing α2-3- and α2-6-linked SIAs. Therefore, only the 9-*O-*acetyl modification on SIA is essential for HCoV-HKU1 glycan binding.

The results demonstrate that glycan recognition is not solely determined by protein structure as N-glycoforms proximal to the Sia-binding pocket influence receptor specificity. It has become increasingly accepted that viral envelope proteins naturally carry immature glycans, specifically at the tip where receptor binding modalities are present [42-44]. Thus, one might argue that expression in GnTI^-^ cells, resulting in Man-5 N-glycans, approximate the receptor binding domain regions. Second, it has been shown that expression of the influenza A virus hemagglutinin in GnTI^-^ cells results in improved SIA binding [31, 45], and this is probably more widely applicable to sialic acid binding proteins, perhaps due to the avoidance of cis-binding [46]. Conclusively, the multivalent presentation of viral glycan binding envelope proteins, that do not contain large complex N-glycosylation, are ideally suited to determine receptor binding properties.

The preferential binding to 9-*O-*acetylated disialic acid raises questions regarding receptor binding and whether the 9-*O-*acetyl α2-3 linked Sia LacNAc ligand would facilitate conformational changes in the spike for S1-CTD exposure [47]. A potential explanation is that HCoV-HKU1 could reversibly interact with low-affinity/high avidity to 9-*O-*acetyl α2-3-linked SIA-containing molecules (mucins), after which the receptor destroying hemagglutinin esterase activity would release the virion from molecules that do not lead to infection. Hereafter, stronger binding to 9-*O-*acetylated disialic acid on glycolipids located on the cell membrane could enable the transitioning of S1-CTD into an open state for proteinaceous receptor binding. Glycolipids are in principle an efficient glycan receptor to enter a cell due to their close proximity to the cell membrane, and it has been shown for SARS-CoV-2 that such structures are important for infectivity [48].

The convergent evolutionary path of HCoV-OC43 and HCoV-HKU1 hemagglutinin esterase activity, and the S1-NTD of BCoV, HCoV-OC43 and HCoV-HKU1 engaging *O-*acetylated SIAs, with preference to certain SIA linkages, indicate host sialoglycome adaptation for optimal replication and tissue tropism. Comparative sialoglycomics of bovids and humans for differences in *O-*acetylation remain to be identified and could enlighten the receptor-binding paradigm of embecoviruses.

## Materials and methods

### Expression plasmid generation

Recombinant HKU1 spike protein NTD (GenBank: DQ339101; AA 14-294) was inserted via Gibson assembly using cDNAs encoding codon-optimized open reading frames of full-length HKU1 spike. HKU1 NTD was cloned into the pcDNA5 expression vector, with a C-terminal human IgG1 Fc, a tobacco etch virus (TEV) cleavage site, a 6xHis-tag, and a Strep-tag (WSHPQFEK; IBA, Germany) [49]. To generate the trimeric HKU1 NTD expression vector, HKU1 NTD cDNA was cloned into the pCD5 expression vector, as previously described [16]. HKU1 NTD was inserted after the signal sequence, in frame with a GCN4 trimerization motif (KQIEDKIEEIESKQKKIENEIARIKK), TEV cleavage site, mOrange2 open reading frame, and a Twin Strep-tag. For the pA-LS NP expression vector [28], domain B of protein A (pA; GenBank: M18264.1, AA 212-270) of *Staphylococcus aureus* and 6,7-dimethyl-8-ribityllumazine synthase (LS; GenBank: AAC06489.1, AA 1-154) of *Aquifex aeolicus* were fused by a Gly-Ser linker and constructed in a pUC57 plasmid by GenScript USA, Inc. The pA-LS sequence was inserted into the pCD5 expression vector, N-terminally fused to a Strep-tag (WSHPQFEK; IBA, Germany).

### Protein expression and purification

Proteins were expressed by transfecting the expression vectors into HEK293T or HEK293S GnTI-cells with polyethyleneimine I (PEI) as previously described [30]. At 6 hours post-transfection, the medium was replaced with 293 SFM II medium (Gibco) supplemented with Primatone 3.0 g/L (Kerry), bicarbonate 3.6 g/L, glucose 2.0 g/L, valproic acid 0.4 g/L, glutaMAX 1% (Gibco), and DMSO 1.5%. Culture supernatants were collected 5 days post-transfection. HKU1 NTD and pA-LS NP protein expression was analyzed using SDS-PAGE and subsequent Western blot on PVDF membrane (Biorad) using StrepMAB-Classic HRP 1:3000 (2-1509-001, IBA). All proteins were purified using Strep-Tactin Sepharose beads (2-1201-002, IBA) as previously described [30].

### Immunofluorescent cell staining

RD cells grown on coverslips were analyzed by immunofluorescent staining. For depletion of surface glycans, cells were incubated for 72 h with 3FNeu5Ac (#5760, Bio-Techne) at 300 µM, Swainsonine (S8195, Sigma) at 10 µM, Ac5GalNTGc [35] at 80 mM, or GENZ-123346 (T4049, TargetMol) at 5 µM. Cells were fixed with 4% paraformaldehyde in PBS for 25 min at RT after which permeabilization was performed using 0.1% Triton in PBS. To present HKU1 NTD Fc chimeras on a NP, NTD Fc proteins were conjugated with pA-LS NP at a 1:1 molar ratio for 1 h at 4 °C. NP-conjugated HKU1 NTD proteins were applied to the coverslips at 50 µg/mL for 1 h at RT. Subsequently, HKU1 NTD trimers or Fc chimeras were precomplexed with primary StrepMAB-Classic HRP (IBA) and secondary goat anti-mouse IgG Alexa Fluor 555 antibody (Invitrogen) applied at a 4:2:1 molar ratio. CTX (C1655, Sigma) was applied at a 1:200 dilution in PBS for 1 h at RT. PHA-L (30280, Vectorlabs), VVL (B-1235, Vectorlabs), and PNA (30272, Vectorlabs) were applied at 10 µg/mL precomplexed with 2.5 µg/mL of streptavidin Alexa Fluor 555 (Thermo Fischer) for 1 h at RT. DAPI (Invitrogen) was used for nuclear staining. Samples were imaged on a Leica DMi8 confocal microscope equipped with a 10× HC PL Apo CS2 objective (NA 0.40). Excitation was achieved with a Diode 405 or white light for excitation of Alexa 555. A pulsed white laser (80 MHz) was used at 549 nm and emissions were obtained in the range of 594–627 nm. Laser powers were 10–20% with a gain of a maximum of 200. LAS Application Suite X was used as well as ImageJ for the addition of the scale bar.

### Glycan microarray

Acetylated structures were printed on glass slides as previously described [10]. HKU1 NTD proteins were either precomplexed with StrepMAB-Classic HRP (IBA) and secondary goat anti-mouse IgG Alexa Fluor 555 (Invitrogen) in a 4:2:1 molar ratio or conjugated with pA-LS NP in a 1:1 molar ratio, in 50 µL phosphate-buffered saline (PBS) with 0.1% Tween-20. In case of pA-LS conjugation, samples were incubated overnight at 4 °C. The mixtures were incubated on the array surface for 90 min. in a humidified chamber. Hereafter, the slides were washed successively with PBS-T, PBS, and deionized water. The slides were dried by centrifugation and scanned as described previously [50]. During processing, the highest and lowest of six replicates were removed and the mean value and standard deviation of the remaining four replicates were calculated. A list of glycans on the microarray is included in table S2.

### Flow cytometry analysis

RD cells were incubated for 72 h with 3FNeu5Ac (#5760, Biotechn) at 300 µM, Swainsonine (S8195, Sigma) at 10 µM, Ac5GalNTGc [35] at 80 mM, GENZ-123346 (T4049, TargetMol) at 5 µM, or a combination of these inhibitors. Approximately 50.000 cells per well were seeded in a 96-well U-bottom plate. HKU1 NTD Fc at 50 μg/mL was conjugated to pA-LS NP at a 1:1 molar ratio for 1 hour on ice and hereafter incubated with the cells for 1 hour on ice. Then, StrepMAB-Classic HRP (IBA) and goat anti-mouse IgG Alexa Fluor 555 (Invitrogen) were added to the cells in a 4:2:1 molar ratio. Trimeric HKU1 NTD and HKU1 NTD Fc at 50 μg/mL were precomplexed with StrepMAB-Classic HRP and goat anti-mouse IgG Alexa Fluor for 1 hour on ice at a molar ratio of 4:2:1. The precomplexed proteins were incubated on the cells for 1 hour at 4°C. CTX (C1655, Sigma) was applied to the cells at a 1:200 dilution. PHA-L (30280, Vectorlabs), VVL (B-1235, Vectorlabs), and PNA (30272, Vectorlabs) at 10 μg/mL were applied with 2.5 μg/mL streptavidin Alexa Fluor 555 (Thermo Fischer) for 1 hour on ice. Viability staining was performed with ViaKrome 808 viability dye (C36628, Beckman Coulter) 1:10,000 diluted in FACS buffer for 5 min at 4°C, followed by centrifugation at 300 rcf for 5 min. After washing with PBS, cells were resuspended in d-PBS supplemented with 0.5% bovine serum albumin (BSA) and 2 mM EDTA. Flow cytometry was performed with the CytoFLEX LX (Beckman Coulter). Data was analyzed using FlowJo software and gated as described in Fig. S2 to select cells and single cells. Mean fluorescence values for triplicates were averaged and standard deviations were calculated.

### Enzyme-linked immunosorbent assay

Nunc MaxiSORP 96-wells plates (Invitrogen) were coated with 10 μg/mL BSM or fetuin in PBS overnight at 4°C, followed by blocking with 3% BSA in 0.1% PBS-T for 3 hours at RT. HKU1 NTD Fc proteins were conjugated with pA-LS NP at a 1:1 molar ratio for 30 min on ice. Trimeric HKU1 NTD, HKU1 NTD Fc, and HKU1 NTD Fc + pA-LS at 50 μg/mL were incubated with StrepMAB-Classic (IBA) and goat αnti-mouse IgG HRP (Invitrogen) at a 4:2:1 molar ratio for 30 min on ice. As positive controls, SNA (B-1305, VectorLabs) and MAL-II (L-1260, VectorLabs) at 50 μg/mL were precomplexed with rabbit anti-biotin HRP (Bethyl) and donkey anti-rabbit Alexa-fluor 555 (Invitrogen) at a 4:2:1 molar ratio. Proteins were added to the plate in a 2-fold serial dilution series and incubated for 1 hour at RT. Ultra TMB-ELISA (34028, Thermo Scientific) was added to the plate and the reaction was stopped after 4 min using 2.5M H_2_SO_4_. Absorbance was measured at 450 nm in a POLARstar Omega plate reader (BMG Labtech).

## Acknowledgements

R.P.dV is a recipient of an ERC Starting Grant from the European Commission (802780). G.J.B is supported by the National Institutes of Health (P41GM103390 and R01HL151617) and by the Netherlands Organization for Scientific Research (NWO TOPPUNT 718.015.003). dr. Lin Liu (CCRCR) and Dr. M.A. Wolfert (Utrecht University) developed, printed and validated the glycan microarray.

## Supplementary Tables

**Table S1.** Coronaviruses and their receptors.

| Genus | Name<sup>a</sup> | Receptor<sup>b</sup> |  |
| --- | --- | --- | --- |
|  |  | S1-NTD | S1-CTD |
| Alphacoronavirus | HCoV-229E | unknown | hAPN [51] |
|  | HCoV-NL63 | unknown | ACE2, heparan sulfate proteoglycans [52, 53] |
| Betacoronavirus | HCoV-HKU1 | 9-O-acetylated sialic acid [15] | TMPRSS2 [54] |
|  | HCoV-OC43 | 9-O-acetylated sialic acid [15, 55] | unknown |
|  | MERS-CoV | Sialoglycans [28, 29] | DPP4 [56] |
|  | SARS-CoV-1 | unknown | ACE2, heparan sulfate proteoglycans [57, 58] |
|  | SARS-CoV-2 | 9-O-acetylated sialic acid [16] | ACE2, heparan sulfate proteoglycans, sialic acids [48, 59, 60] |

**Notes:** <sup>a</sup>HCoV, human coronavirus; MERS-CoV, Middle East respiratory syndrome coronavirus; SARS-CoV, severe acute respiratory syndrome coronavirus. <sup>b</sup>hAPN, human aminopeptidase N; ACE2, angiotensin-converting enzyme 2; TMPRSS2, transmembrane serine protease 2; DPP4, dipeptidylpeptidase 4

**Table S2.**
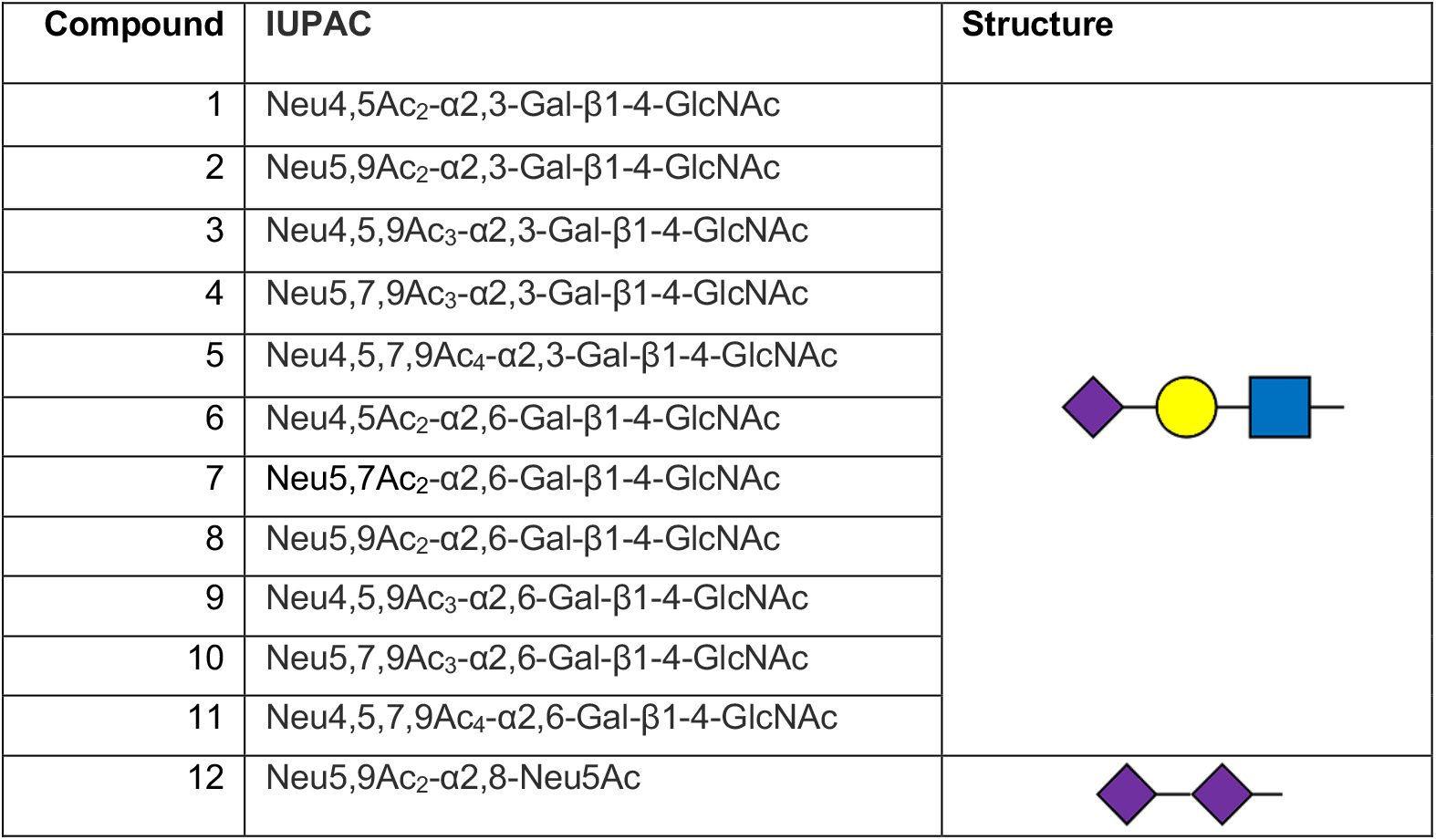
Glycans imprinted on the micro array.

| Compound | IUPAC | Structure |
| --- | --- | --- |
| 1 | Neu4,5Ac<sub>2</sub>-α2,3-Gal-β1-4-GlcNAc |  |
| 2 | Neu5,9Ac<sub>2</sub>-α2,3-Gal-β1-4-GlcNAc |  |
| 3 | Neu4,5,9Ac<sub>3</sub>-α2,3-Gal-β1-4-GlcNAc |  |
| 4 | Neu5,7,9Ac<sub>3</sub>-α2,3-Gal-β1-4-GlcNAc |  |
| 5 | Neu4,5,7,9Ac<sub>4</sub>-α2,3-Gal-β1-4-GlcNAc |  |
| 6 | Neu4,5Ac<sub>2</sub>-α2,6-Gal-β1-4-GlcNAc |  |
| 7 | Neu5,7Ac<sub>2</sub>-α2,6-Gal-β1-4-GlcNAc |  |
| 8 | Neu5,9Ac<sub>2</sub>-α2,6-Gal-β1-4-GlcNAc |  |
| 9 | Neu4,5,9Ac<sub>3</sub>-α2,6-Gal-β1-4-GlcNAc |  |
| 10 | Neu5,7,9Ac<sub>3</sub>-α2,6-Gal-β1-4-GlcNAc |  |
| 11 | Neu4,5,7,9Ac<sub>4</sub>-α2,6-Gal-β1-4-GlcNAc |  |
| 12 | Neu5,9Ac<sub>2</sub>-α2,8-Neu5Ac |  |

**Table S3.** EC50 values of HKU1 NTD Fc binding to BSM. EC50 values were based on BSM ELISA data (Figure 5A). A significant difference in HEK293-derived HKU1 NTD Fc + pA-LS binding to BSM was observed in comparison to HEK293-derived HKU1 NTD Fc.

| Protein | EC50 | p-value |
| --- | --- | --- |
| Fc, HEK293T | 19.064 | <0.0001 |
| Fc + pA-LS, HEK293T | 9.262 |  |
| Fc, GnTI | 7.819 | n.s. |
| Fc + pA-LS, GnTI | 6.786 |  |

## Supplementary figures

**Fig. S1.**
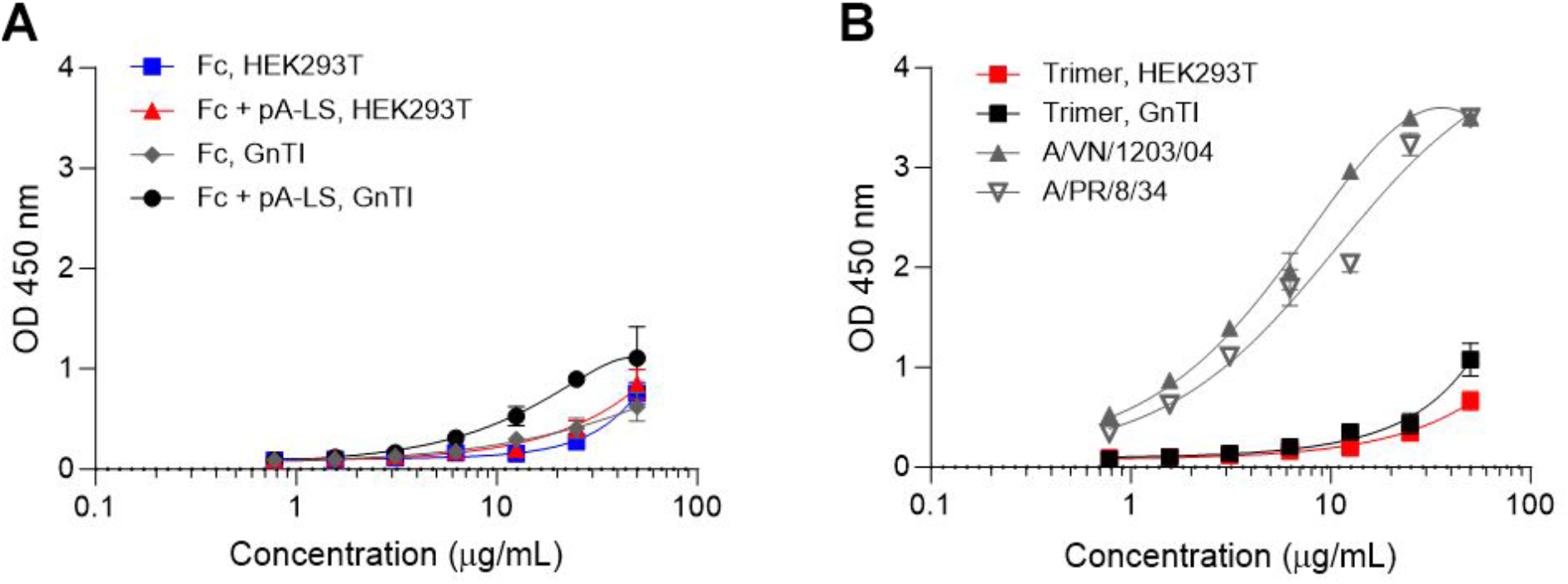
O-acetylation dependent virolectin binding. **(A)** HKU1 NTD Fc and HKU1 NTD Fc + pA-LS (GnTI-derived and 293-derived) did not display binding to fetuin. **(B)** Presence of sialic acids on fetuin was confirmed by using A/Vietnam/1203/2004 (A/VN/1203/04) and A/Puerto Rico/8/1934 (A/PR/8/34). Trimeric HKU1 NTD (GnTI-derived and 293-derived) did not bind fetuin.

**Fig. S2.**
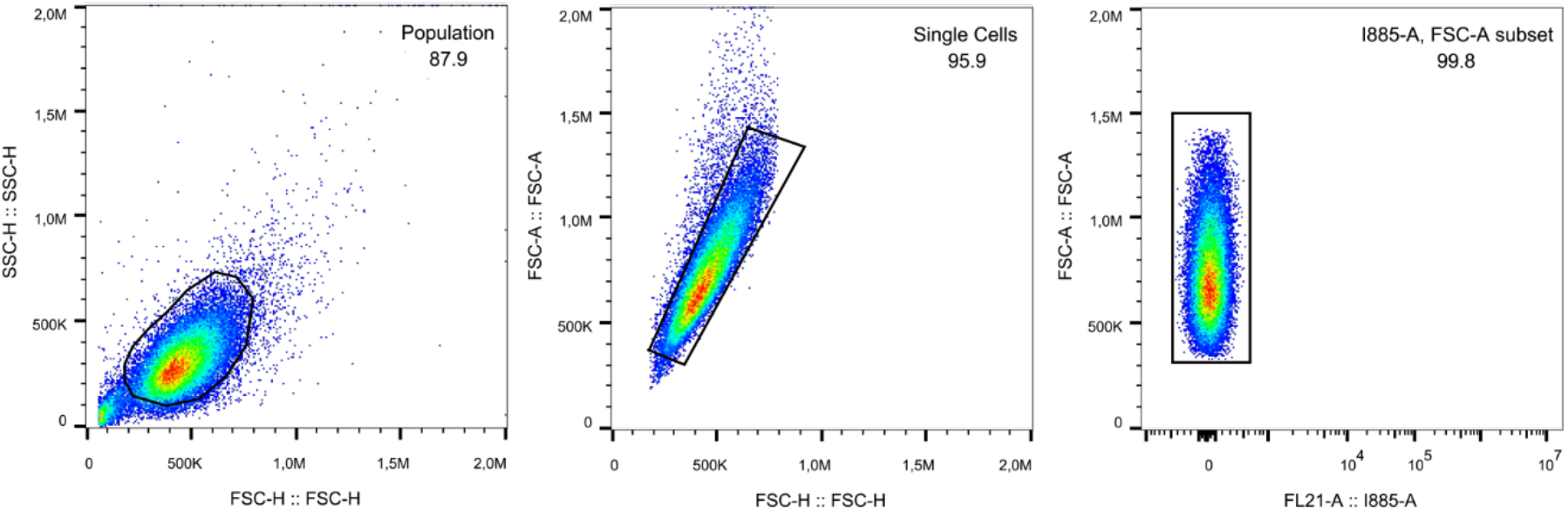
Gating strategies for flow cytometry. Gating was performed to select for cell population, followed by singlets and then live-cells using ViaKrome 808 viability dye.

## Notes

### Competing Interest Statement

The authors have declared no competing interest.

